# Genotypic evidence of vector-mediated selection on *Plasmodium falciparum* in infected field-collected *Anopheles* mosquitoes

**DOI:** 10.64898/2026.10.23.759723

**Authors:** Ailsa Craven, Ousmane Akone-Ella, Katrina A. Kelley, Theo Mouillaud, Sandrine Eveline Nsango, Sarah Perkins, Angela M. Early, Diego Ayala, Daniel E. Neafsey

## Abstract

Malaria-causing *Plasmodium* parasites are transmitted between vertebrate hosts via diverse species of *Anopheles* mosquitoes. The P47 protein in *Plasmodium falciparum* has been shown to affect transmission in various anopheline species by mediating evasion of mosquito innate immunity in an allele-specific manner. The *Pfs47* gene that encodes P47 exhibits high genetic divergence across continents, as well as high genetic diversity within many parasite populations, but the impact of polymorphisms on transmission success in different sympatric vectors has not been previously studied in a field context. Here we report parasite genotyping data from naturally infected, field-collected mosquitoes at sites harboring at least two prevalent sympatric vector species, targeting *Pfs47* in addition to the *CSP*, *AMA1*, *SERA2*, and *TRAP* parasite antigens. Collections took place at Ndangueng, Cameroon and Benguia, Gabon between November 2019 and July 2021, yielding infected *Anopheles gambiae* at both sites in addition to *Anopheles moucheti* in Cameroon and *Anopheles funestus* in Gabon. We found no significant difference in the mean number of parasite strains in infected sympatric mosquito species from each site. However, *Pfs47* exhibited a significant difference in the prevalence of an amino acid polymorphism (I248L) between *An. moucheti* and *An. gambiae* in Cameroon. Polymorphisms at two additional positions (N272Y/I and P369H) displayed significant linkage disequilibrium with I248L, and there were significant differences in the distribution of *Pfs47* haplotypes between sympatric mosquito species at both sites. These findings demonstrate divergent selection on *P. falciparum* mediated by sympatric vector species, suggesting that parasite genotype significantly impacts local vectorial capacity by modulating species-specific vector competence.

## Introduction

Malaria remains an important threat to global public health, with an estimated 282 million cases and 610,000 deaths in 2024.^1^ Human malaria-causing *Plasmodium* parasites are transmitted between hosts via the blood-feeding behavior of *Anopheles* mosquitoes. However, parasites ingested during a blood meal from one infected host are not simply re-injected into a subsequent feeding host. Rather, parasites must undergo elaborate biological changes within the mosquito over a period of at least 7-10 days, including completion of the compulsory sexual phase of the parasite lifecycle and development from a zygote into the infectious sporozoite life stage, while evading the mechanisms of mosquito innate immunity mediated by over 200 genes belonging to 18 gene families.^2^ Vectorial competence, the inherent ability of a mosquito to transmit the malaria parasite, is therefore an important determinant of local vectorial capacity, here defined as the efficiency with which the local assemblage of one or more mosquito species transmits malaria parasites. This capacity is also determined by factors such as vector population density and biting rate.^3^ Variability among species in vectorial competence is extensive, of the over 450 described *Anopheles* species, 70 are demonstrated competent vectors and around 50 are classified as dominant vector species.^4,5^ The evolutionary history of association between *Plasmodium* parasites and *Anopheles* mosquitoes is therefore important to study for understanding the contemporary global distribution of malarial disease as a function of vectorial competence.

The most thoroughly characterized parasite protein associated with vectorial competence is P47 (gene *Pfs47*), a member of the 6-cysteine protein family that disrupts mosquito midgut immune signaling when expressed in early mosquito-infective stages of the parasite (female gametocytes and ookinetes).^6^ Genetic knockouts of *Pfs47* and its ortholog *Pbs47* in the mouse parasite *Plasmodium berghei* demonstrate comprehensive effects on parasite survival in immune-competent mosquitoes.^7,8^ Further, experiments using laboratory-adapted lines of *P. falciparum* from Africa, Asia, and South America demonstrated that parasites exhibit enhanced survival in *Anopheles* species native to the same continent as the parasite lines (Africa: *Anopheles gambiae*; Asia: *Anopheles dirus*; South America: *Anopheles albimanus*), leading to the ‘lock and key’ hypothesis of parasite-vector coevolution, which holds that differences among parasites in *Pfs47* sequence (the ‘key’) are driven by population or species-level differences in the vector receptor to *Pfs47* (P47Rec, the ‘lock’),^9^ as molecular mismatch results in parasite detection and melanization via the JNK signaling pathway.^10^

Parasite population genomic studies support the hypothesis of divergent selection pressures by different vector species. *Pfs47* was observed to exhibit the highest divergence level of any gene between continental populations of *P. falciparum* in the first global genome-wide survey of diversity in that species,^11^ and the *Pfs47* ortholog in *Plasmodium vivax* (*Pvs47*) similarly exhibited very high divergence between parasite populations in the Americas vs. Asia/Africa.^12^ In the Americas, *Plasmodium* parasites are primarily transmitted by anopheline vectors belonging to the *Nyssorhynchus* subgenus, which is estimated to have diverged from Old World subgenera 30-100 million years ago.^13–15^ When *Plasmodium* parasites were brought to the Americas from Africa and Europe during the colonial era, it is hypothesized that they were subject to strong selection pressure by anophelines that were highly evolutionarily divergent from their Old World counterparts, leading to the divergence observed at *Pfs47* and other loci associated with vector adaptation.^16–18^

In addition to divergence between allopatric parasite populations, *Pfs47* frequently exhibits genetic diversity within populations of *P. falciparum* and *P. vivax*. In many settings vectorial capacity is determined by more than one sympatric anopheline vector species, leading to the hypothesis that adaptation for optimized transmission by different sympatric anopheline species leads to balancing selection on *Pfs47* in local parasite populations.^19^ Testing this hypothesis directly by genotyping parasites within infected mosquitoes has proven challenging for technical reasons, as many vector species are difficult to colonize, and the prevalence of infected mosquitoes is low in most settings.^20^ While no systematic effort has been made to associate *Pfs47* diversity with local vectorial taxonomic diversity at a global level, recent work has led to better characterization of *Pfs47* variation in many settings, and an association between local *Pfs47* diversity and the presence of several anopheline species has been observed in Africa.^21^

In this manuscript we address the gap in direct evidence for the role of vector-mediated selection in the maintenance of diversity at *Pfs47* and several other parasite loci at a population level. We performed extensive collection and genotyping of infected mosquitoes at two high transmission sites in Central Africa: Ndangueng, Cameroon and Benguia, Gabon. Both of these sites exhibited a high diversity of sympatric vector species as well as high parasite genetic diversity, enabling us to collect a large number of naturally-infected mosquitoes at each site and search for associations between parasite genotype and mosquito taxon. We report direct evidence that sympatric malaria vectors exert taxon-specific selection on parasites as a function of their genotype, suggesting that parasites in settings with high vector diversity may be subject to divergent selection pressures for transmission, with possible negative consequences for overall local vectorial capacity.

## Results

### Mosquito Collection

Collection of *Anopheles spp.* mosquitoes took place by live capture across four collection missions in two different sites: Ndangueng, Cameroon and Benguia, Gabon (Fig 1A). Overall, 7,414 mosquitoes were collected (3,866 in Cameroon and 3,548 in Gabon) and 4.57% (339) of the mosquitoes collected were positive for *P. falciparum* by qPCR (Supplementary Table 1). Five species of collected *Anopheles* were positive for *P. falciparum* across the two sites (Table 1). Among them, *Anopheles hancocki* in Cameroon and *Anopheles paludis* in Gabon, only had one infected sample and were therefore not included in analyses. Infected *An. gambiae* were present at both sites and coincidentally exhibited a similar infection prevalence of 6.4% (86 out of 1352 in Cameroon and 138 out of 2143 in Gabon). In addition to *An. gambiae*, there were 108 infected *An. moucheti* samples collected from Cameroon and 21 infected *An. funestus* samples collected in Gabon (Fig 1B/C). Both *An. moucheti* and *An. funestus* had lower *P. falciparum* infection prevalence than *An. gambiae*, with 4.1% (83 out of 2041 in Cameroon) and 4.8% (17 out of 356 in Gabon) of mosquitoes testing positive for *P. falciparum*, respectively. The second most common species of mosquito caught in Gabon was *An. paludis* (n=550), but only one tested positive for *P. falciparum*, making the infection prevalence for *An. paludis* in Gabon approximately 0.2%. Both *An. moucheti* and *An. funestus* were collected at both sites, however each of these species exhibited *P. falciparum* infection at only one site (Table 1, Supplementary Table 1).

**Figure 1.**
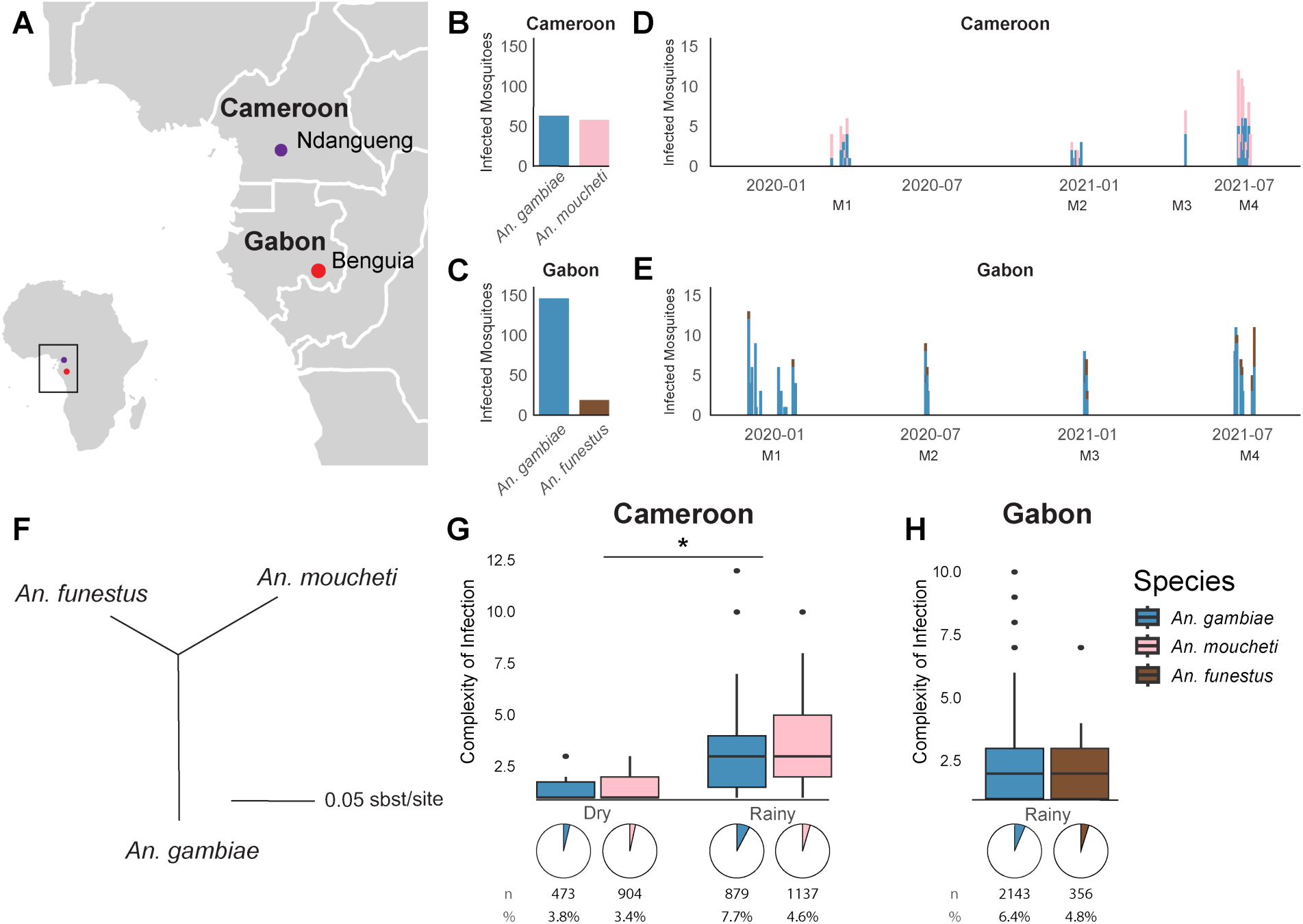
Spatial and temporal distribution of infected *Anopheles spp*. A) Samples were collected at Ndangueng in Cameroon and Benguia in Gabon. In Cameroon (B) a total of n=63 *An. gambiae* and n=58 *An. moucheti* were collected and in Gabon (C) a total of n=146 *An. gambiae* and n=19 *An. funestus* were collected. Collections in Cameroon (D) and Gabon (E) occurred via four missions at each site spanning November 2019 to August 2021. F) An unrooted tree generated from the complete mitogenomes of each *Anopheles* species using IQTree.^55^ G) Complexity of Infection was calculated using 4CAST amplicon sequencing in both Cameroon and H) Gabon. Total mosquitoes collected (n) and infection prevalence (%) are listed along with prevalence charts by season for each population. Comparisons of COI between species and seasons was performed by a pairwise Wilcoxon test.

**Table 1.** Infected Mosquito Collection Summary.

| Country | *Anopheles*<br>Species | Mission | Collected | *P. falciparum*<br>infected | *P. falciparum*<br>Prevalence |
| --- | --- | --- | --- | --- | --- |
| Cameroon | *An. gambiae* | M1 | 473 | 18 | **0.064** |
|  |  | M2 | 489 | 26 |  |
|  |  | M3 | 123 | 0 |  |
|  |  | M4 | 267 | 42 |  |
|  | *An. hancocki* | M2 | 5 | 1 | **0.200** |
|  | *An. moucheti* | M1 | 904 | 31 | **0.041** |
|  |  | M2 | 256 | 5 |  |
|  |  | M3 | 54 | 3 |  |
|  |  | M4 | 827 | 44 |  |
| Gabon | *An. funestus* | M1 | 101 | 2 | **0.048** |
|  |  | M2 | 48 | 1 |  |
|  |  | M3 | 56 | 4 |  |
|  |  | M4 | 151 | 10 |  |
|  | *An. gambiae* | M1 | 1271 | 56 | **0.064** |
|  |  | M2 | 303 | 11 |  |
|  |  | M3 | 196 | 19 |  |
|  |  | M4 | 373 | 52 |  |
|  | *An. paludis* | M1 | 68 | 0 | **0.002** |
|  |  | M2 | 191 | 0 |  |
|  |  | M4 | 291 | 1 |  |
|  | sp | M1 | 140 | 8 | **0.055** |
|  |  | M2 | 20 | 0 |  |
|  |  | M3 | 40 | 3 |  |
|  |  | M4 | 37 | 2 |  |

Number of total and *P. falciparum*-infected mosquitoes collected by species. Only *Anopheles* species with a *P. falciparum* prevalence greater than 0 are shown.

In Cameroon, approximately equal numbers of infected mosquitoes were collected between species across the four missions (Fig 1D, p= 0.80 by Fisher’s Exact Test). In Gabon, there were fewer infected *An. funestus* collected across all the missions, and the relative number of infected *An*. *funestus* compared to *An*. *gambiae* increased moderately across the four missions in a non-statistically significant manner (Fig 1E, p = 0.067 by Fisher’s Exact Test).

The mitochondrial genomes of *An*. *gambiae, An. moucheti,* and *An*. *funestus* were aligned and used to create an unrooted Maximum Likelihood tree to approximate the relative evolutionary distances among the focal vector species in this study (Fig 1F). *An. moucheti* and *An*. *funestus* were the closest pair, with a divergence of 0.048 nucleotide substitutions (subs)/site; *An. moucheti* and *An. gambiae* were the most distant exhibiting a rate of 0.102 subs/site, and *An. funestus* and *An*. *gambiae* exhibited a divergence of 0.071 subs/site.

### Amplicon sequencing of highly diverse loci reveals no differences in COI or allele distribution between sympatric populations

A target amplicon panel (4CAST) of highly diverse loci was used to calculate complexity of infection (COI) (the number of genetically distinct parasite clones) for each infected mosquito sample. There were no significant differences in COI between sympatric mosquito populations at either site (Cameroon *gambiae* COI: 3.28, *moucheti* COI: 3.24, p = 0.9686; Gabon *gambiae* COI: 2.73, *funestus* COI: 2.25, p= 0.4432; comparisons done by pairwise Wilcoxon test; Fig 1G,H). In Cameroon, where mosquitoes were collected in both dry and rainy seasons, there was an increase in COI in samples collected during rainy seasons relative to the dry seasons (COI = 3.05 vs. 1.53, respectively; Fig 1G,H, rainy-dry *W* = 2301.5, *p* = 0.003958).

Parasite antigen variants in 4CAST loci were compared between vector species and differences in amino acid variant prevalence between species were tested for statistical significance within each location via Fisher’s Exact Test (Table 2, Supplementary Figure 1). There were no significant associations between 4CAST antigen SNPs and vector species in either location after correcting for multiple testing (Bonferroni-adjusted p > 0.05).

**Table 2.** 4CAST Loci comparisons by Fisher’s Test.

|  |  | **p value** | **adj. p value** |
| --- | --- | --- | --- |
| **CSP** | Cameroon | 0.362 | 1.000 |
|  | Gabon | 0.679 | 1.000 |
| **AMA1** | Cameroon | 0.402 | 1.000 |
|  | Gabon | 0.466 | 1.000 |
| **SERA2** | Cameroon | 0.928 | 1.000 |
|  | Gabon | 0.063 | 0.508 |
| **TRAP** | Cameroon | 0.241 | 1.000 |
|  | Gabon | 0.946 | 1.000 |

Each locus at each location was compared by Fisher's Exact Test for Count Data with simulated p-value (based on 2000 replicates). P values were adjusted for multiple testing using Bonferroni correction.

### *Pfs47* displays position-specific prevalence differences between sympatric vector species

As parasite whole genome sequencing did not yield sufficient coverage to perform genome-wide scans for variants differentially distributed between sympatric vector samples, we conducted a targeted analysis of *Pfs47* genetic variation given prior evidence of mosquito-driven selection (Fig 2A).^8,9,22^ Publicly available *P. falciparum* sequencing data from West African samples showed four variants with frequencies greater than 10%: residues 194, 248, 272, and 369. One of the residues (272) is polyallelic with SNPs at nucleotide positions 814 and 815 causing different nonsynonymous mutations. The primers successfully amplified a region within the coding sequence extending from residue 224 to 390, covering three out of four variant positions.

**Figure 2.**
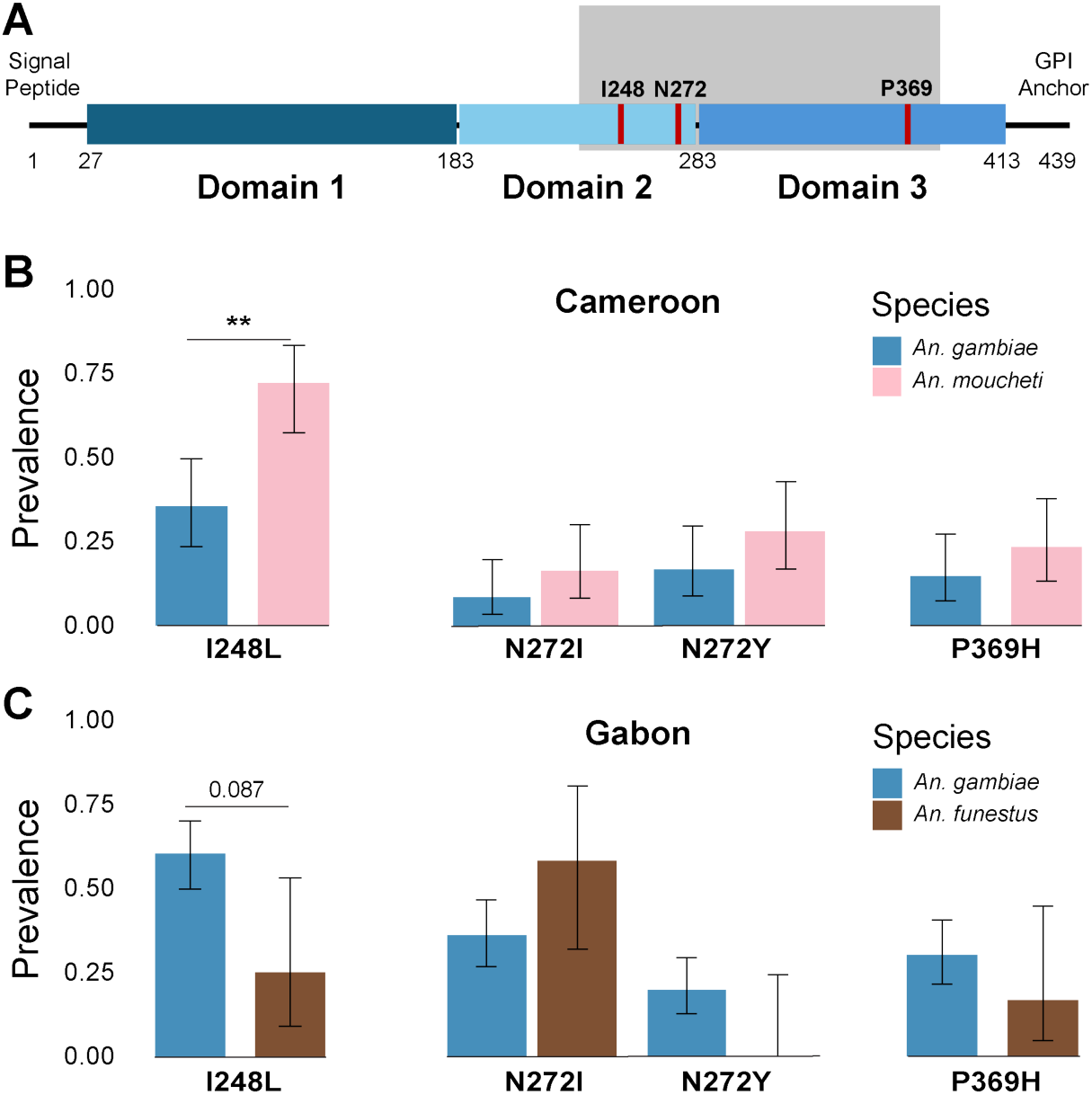
Sympatric mosquito populations display differences in I248L prevalence. A) A representative diagram of *Pfs47* domains. The region of the gene that was amplified and sequenced is shown by the gray box. The three positions that had two or more SNPs present with a minor allele prevalence above 5% are marked in red and labelled. Prevalence of the non-reference alleles in populations of sympatric mosquitoes in Cameroon (B) and Gabon (C). Comparisons of prevalence between species was performed by Fisher’s Exact test and p values were adjusted for multiple comparisons using Bonferroni correction. (**p < 0.01). All error bars show 95% Wilson confidence intervals.

The four most common variants identified in publicly available data - I248L, N272I, N272Y, and P369H - were also the four most common amino acid variants in our data (Supplementary Figure 2) with one of the six other variants appearing in two samples and the remaining five only appearing in one. All further analyses were done with the four SNPs that had a prevalence above 5%.

Each polymorphic position was examined for differences in variant prevalence between vector species at both sites (Fig 2). There was a significant difference in the prevalence of I248L between *An*. *gambiae* and *An*. *moucheti* in Cameroon (I248L Prev in *An. gambiae* = 0.35, *An. moucheti* = 0.72, adj. p = 0.0021, Fig 2B). Due to the small sample size, the difference in I248L prevalence between *An. gambiae* and *An. funestus* was not significant after correction for multiple comparisons (I248L Prev in *An. Gambiae* = 0.60, *An. funestus* = 0.25, adj. p = 0.087, Fig 2C). There was no significant difference in variant prevalence between sympatric species at other positions at either site (Cameroon: *gambiae-moucheti* N272 adj. p = 0.22, P369 adj. p = 0.42; Gabon: *gambiae-funestus* N272 adj. p = 0.32, P369 adj. p = 0.50, comparisons done by Fisher’s Exact test with Bonferroni correction, Fig 2B); given that their overall prevalence is lower than I248L it is possible that there could be a significant difference if the sample size were larger. These results accord with previous studies, showing I248L as a position of interest in experimental comparisons of 3D7 and 7G8 strains of *P. falciparum*.^22^

### Differences in I248L prevalence among vector species are not due to micro-temporal stratification

Because the mosquitoes were collected over the course of 16-18 months, it is possible that observed differences in parasite SNP frequencies are a consequence of temporal changes in frequency rather than selection by vector species. To exclude this alternative hypothesis, the prevalence of I248L at both sites across all missions was compared. There was no significant difference in overall prevalence of I248L across missions (Fig 3A p= 0.80, 3B p= 0.82). The difference in I248L prevalence between *An. gambiae* and *An. moucheti* in Cameroon remained consistent across missions (Fig 3C), and Mission 4 had a large enough sample size to generate a statistically significant difference in prevalence. To further confirm these findings, a logistic regression model was run with I248L as the outcome, mosquito species as the explanatory variable, and date collected as a potential confounding variable (Supplementary Figure 3). After adjusting for date, the predictive power of species on the probability of I248L presence did not change and remained significant in Cameroon. Similarly in Gabon, adjusting for date did not alter the predictive power of species on the probability of I248L presence.

**Figure 3.**
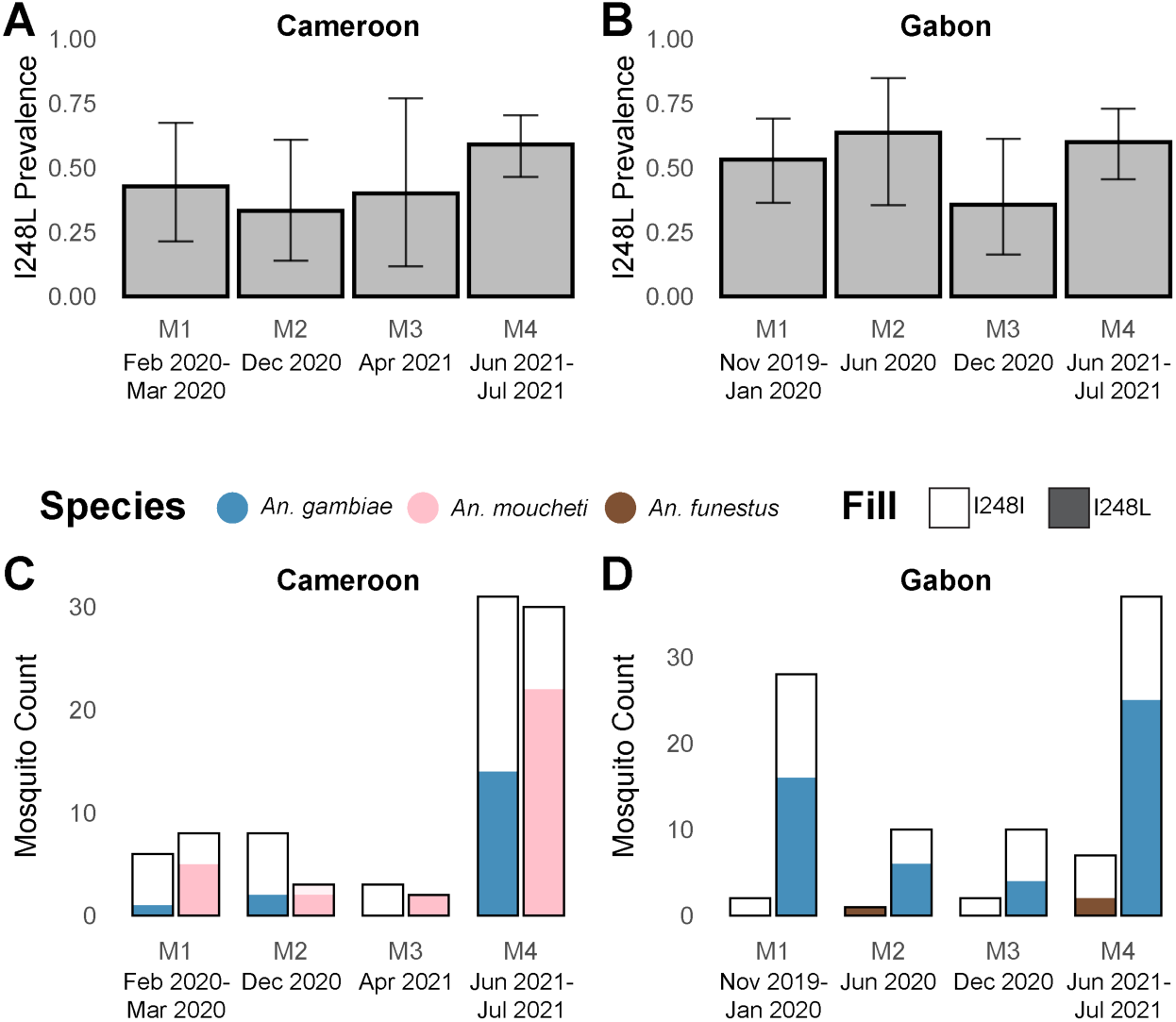
I248L prevalence does not change over time and species-specific differences in prevalence remain consistent across missions. The overall prevalence of the I248L SNP in Cameroon (A) and Gabon (B) across each mission. All error bars show 95% Wilson confidence intervals. Count of infected mosquitoes separated by species that have either I248I or the I248L allele across each mission in Cameroon (C)and Gabon (D).

### *Pfs47* displays differences in haplotype distribution and linkage disequilibrium between sympatric vector species

We explored linkage disequilibrium among *Pfs47* variants in Gabon and Cameroon. To conduct haplotype-level analyses, samples with heterozygosity at more than one position were excluded (n=15) as the genotypes cannot be phased. We describe haplotypes as a trio of letters corresponding to the nucleotides at each position (positions 248, 272, and 369; Figure 4). Of the twelve possible haplotypes that could exist, IIH, IYP, and IYH were not found among these samples, and some alleles at positions 272 and 369 were only observed in a certain background of 248.

**Figure 4.**
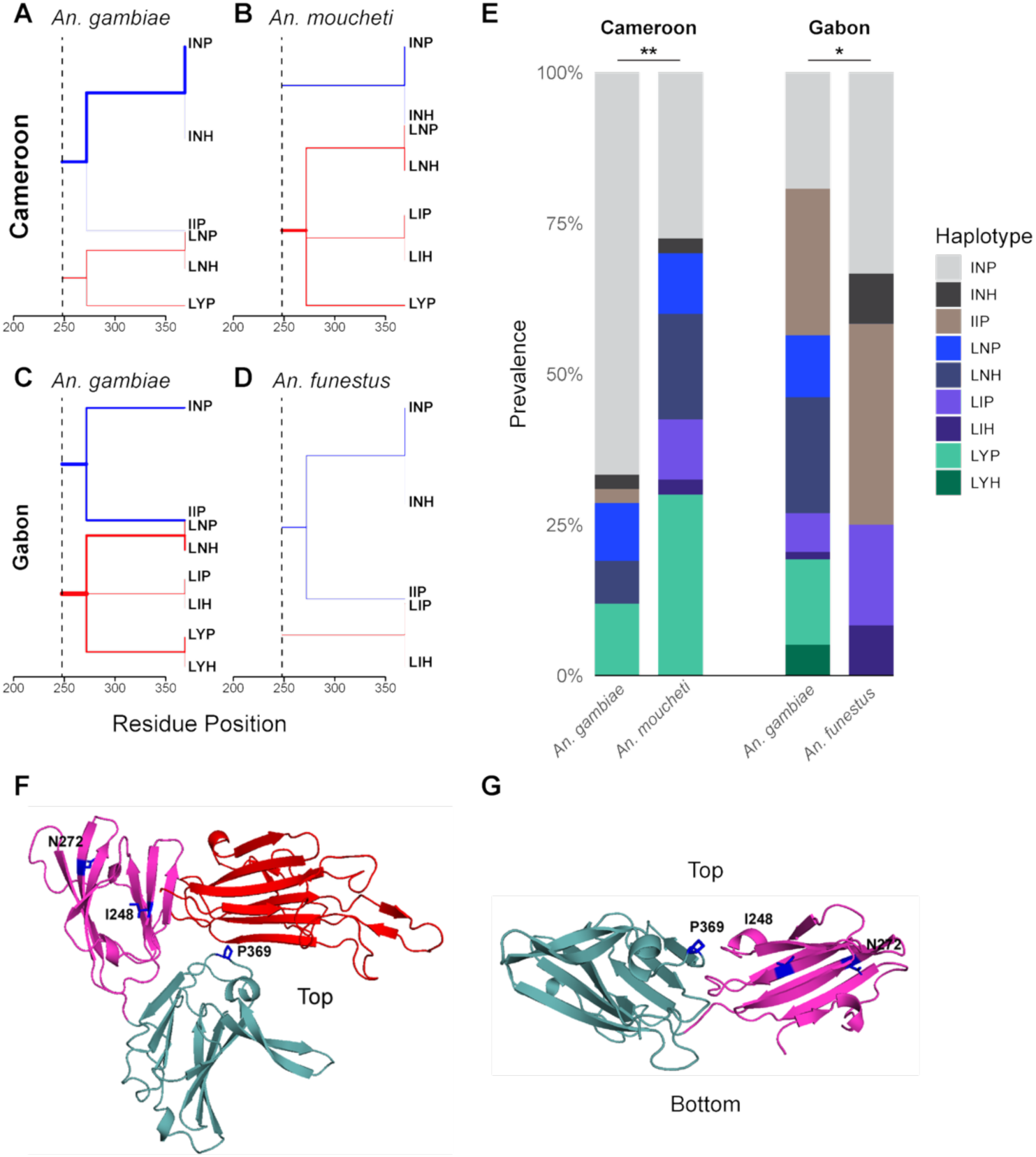
Sympatric mosquito populations display differences in the distribution of *Pfs47* haplotypes. A-D) Bifurcation diagrams for each mosquito population anchored on I248. Relative prevalence of each haplotype defined by SNPs at amino acid positions 248, 272, and 369 is indicated by line weight. Blue indicates I248I and red indicates I248L. E) Distribution of the prevalence of haplotypes, as represented by the residue present at each of the three positions of interest, for each mosquito population. Comparisons of haplotype distributions were carried out by Fisher’s Exact Test. F/G) AlphafoldDB prediction of 3D7 P47 excluding the signal peptide (1-27) and the GPI anchor (413-439). Residues of interest are highlighted in dark blue and labelled. F) View from the “top” a.k.a. the side opposite the GPI anchor. G) View from the side with domain one removed. (*p < 0.05, **p < 0.01).

To determine if there were population-specific differences in linkage disequilibrium, bifurcation diagrams were plotted around I248L for each population (Fig 4A-D) and pairwise correlation (R^2^ values) was calculated for N272I, N272Y, and P369H against I248L (Supplementary Figure 4). The bifurcation diagrams show a complete absence of IYX haplotypes, and a preference for LYX over LIX across mosquito populations. There was also a difference in the distribution of R^2^ values across positions and samples. We hypothesized that there would be differences in the distribution of these haplotypes between sympatric species. There was a statistically significant difference in the distribution of haplotypes between sympatric species at both collection sites (*gambiae-moucheti* p = 0.003; *gambiae-funestus* p = 0.032 by Fisher’s Exact Test, Fig 4E).

We hypothesized that these three positions might lie close to each other in the tertiary structure of P47 given the observed linkage disequilibrium. Both I248 and N272 are in the immunogenic Domain 2, however P369 is in Domain 3. Examination of residue positions using a high-confidence Alpha Fold-generated prediction of P47 structure shows that the positions are not located proximally to each other, however all three positions are closer to the outward face (away from the GPI anchor) of P47 (Fig 4F,G).

## Discussion

In order to be transmitted from person to person, *Plasmodium* parasites must invade mosquito midguts and salivary glands while evading complement activation and other attacks from the mosquito immune system. The effects of mosquito immunity, and other non-immune determinants of infection, are well characterized in laboratory models, but how these effects differ across the multitude of vector-competent *Anopheles* species is less explored. Vector-competent *Anopheles* often have large overlaps in range; many areas have multiple malaria vector species with varying levels of transmission competency^23^ so parasites may be subject to selective pressures from multiple highly genetically divergent vectors simultaneously.

Here we present the first direct investigation of local vector-mediated selection on *P. falciparum* using over 7,000 field-caught mosquitoes from Gabon and Cameroon. Initial qPCR revealed heterogeneity in infection prevalence across mosquito species. Although *An. paludis* was the second most prevalent species found overall, parasite infection was rare (0% positive in Cameroon and 0.2% positive in Gabon, Supplementary Table 1), despite other studies demonstrating sporozoite prevalence from 1-10% in other regions of Sub-Saharan Africa.^24–26^ This may be the result of high regional specificity of *P. falciparum* to *An*. *paludis,* the zoophilicity^27^ and exophilicity^28^ of this species, or of conflation of *An. paludis* with a similar species, as the study which used molecular methods to confirm taxonomy estimated sporozoite prevalence around 1%.^25^ The circumsporozoite rate of *An. paludis* captured in Cameroon between 1998-2003 was similarly low, at approximately 1%.^27^ Polyclonality estimates using the 4CAST amplicon panel revealed an average COI of 2.7 in Gabon and 3.6 in Cameroon during rainy seasons, slightly lower than estimates from other studies.^29,30^ *A. gambiae* COI closely reflected overall COI across both sites (3.5 in Cameroon and 2.7 in Gabon) which could reflect differences in malaria prevalence between sites or differences in vector competence between these geographically distinct populations.^31^ Additionally, there were lower overall COI values in the dry season (Fig. 1GH), consistent with the decrease in mosquito prevalence and transmission during those months.

While human host immunity is thought to select for genetic diversity in parasite antigens,^32–34^ the extent to which vector-mediated selection contributes to antigenic diversity is unclear. Previous studies have isolated parasite-vector interactions involving parasite antigens notable in the acquired immune responses of hosts, such as that proposed between the antigen *TRAP* and mosquito salivary gland protein *Saglin*.^35^ We observed no significant associations between vector species and amino acid polymorphisms in *CSP*, *AMA1*, *SERA2*, and *TRAP*. Similarly, despite evidence that mosquito selective pressures may be driving CSP genetic structure in *P. vivax*,^36^ there were no observed differences in *CSP* SNPs corresponding to vector species or location. While this survey was modest in terms of the number of loci and vector species screened, these results do not provide support for a strong contribution of mosquito vectors to ongoing balancing selection on parasite antigens, suggesting human-host-mediated selection is the primary driver of antigen diversity.

We also applied capillary sequencing to determine whether there were differences in *Pfs47* SNP prevalence based on vector species or collection location. There were species-specific differences in I248L residue prevalence between *An. gambiae* and *An. moucheti* in Cameroon, but there were no other significant differences in SNP prevalence between species or locations, possibly due to small sample size or low SNP frequency (Fig 2). There were no confounding effects of collection date on these species-specific differences in prevalence (Fig 3). There was also evidence of linkage disequilibrium, as notably N272Y is only seen in the background of I248L (Fig 4AD), and there were significant differences in the distribution of haplotypes across sympatric populations at both sites (Fig 4E). The distributed locations of these amino acid residues as predicted by Alphafold suggest that the *Pfs47* protein may be evolving broadly in response to selective pressure from native mosquito species (Fig 4FG).

Previous work has demonstrated that *Pfs47* is highly geographically structured at a continental scale, and has been implicated in parasite evasion of the *Anopheles* immune response.^8,37^ SNPs, particularly at the 707, 725, and 740 nucleotide position, are highly predictive of continent-of-origin of *Pf*^38^ and, along with 742, separate the 3D7 and 7G8 haplotypes that have been extensively studied in laboratories.^22^ Sub-continental studies have demonstrated high minor allele frequencies at different positions, notably nucleotide position 81 (E27D) has a high minor allele frequency in Kenya.^39,40^ When examining population genomic data from West African human samples, the nucleotide positions 742 (I248L), 814 (N272Y), 815 (N272I), and 1106 (P369H) have the highest minor allele frequencies, with 581 (P194H) following as it appears close to fixation.

The large difference in I248L prevalence between sympatric vector species (37% n=0.35/0.72 *gambiae/moucheti* in Cameroon and 35% n=0.60/0.25 *gambiae/funestus* in Gabon) suggests that strong mosquito-driven selection is occurring. Given this prevalence difference, we would expect up to a third of parasites to be removed via immune-mediated clearance due to incompatible *Pfs47* alleles. Both sites display differences in *P. falciparum* prevalence between sympatric species (36.1% drop in prevalence between *An. gambiae* and *An. moucheti* n=0.064/0.041; 25.8% drop in prevalence between *An. gambiae* and *An. funestus* n=0.064/0.048), however, exact estimations of a selection coefficient are hampered by a large number of haplotypes, and high COI of both human and mosquito infections. Additionally, P47 is expressed in female gametes^7,8^ so only one of the two P47 alleles is present after fertilization which could allow for the transmission of “incompatible” *Pfs47* alleles silently contributed by male gametes.

This study had some limitations. Despite the thousands of mosquitoes collected, there were still relatively small numbers of infected specimens obtained for *An. funestus*, limiting the statistical comparisons that could be made. This study also does not detect all loci involved in divergent local selection by sympatric vector species as it was not possible to perform whole genome sequencing. Because we used capillary sequencing for *Pfs47*, we could only calculate prevalence and not frequency of both SNPs and haplotypes, which means we are likely underestimating the SNP association between sympatric species.

Despite these shortcomings, our results provide the first strong evidence of divergent selection on *P. falciparum* parasite genotype by sympatric vector species. Differences in both SNPs and haplotype frequencies of *Pfs47* in parasites infecting sympatric *Anopheles* species provide evidence for divergent selective pressure on parasites in areas with multiple competent vectors. While *Pfs47* is not the only *Plasmodium* locus implicated in mediating parasite transmission,^41^ it remains an important and useful benchmark locus for gaining a greater understanding of how vector species and location shape parasite evolution. Further work is needed to enhance understanding of mosquito immune mechanisms and their role in generating local diversity and geographic divergence in parasite populations.

Malaria control efforts such as gene drive-mediated population replacement must be tailored to the location of interest to be effective.^42,43^ Different mosquito species have varying levels of epidemiological importance due to parasite compatibility, feeding behavior, and other traits that determine vectorial capacity, which may vary by region.^44,45^ As we gain more detailed surveys of vector abundance and infection prevalence, it is becoming clear that mosquito species that were classified as secondary vectors such as *An. moucheti* are contributing more to the spread of malaria than previously assumed and have been considered a co-primary vector in certain regions^23,46^ The vectorial capacity of a region is a function of local vector diversity and competence, but our work demonstrates that local parasite genetic background may also influence vector competence in a manner that varies by vector species, indicating that the vectorial capacity in a location is not solely defined by the presence and density of all possible vector species. Our work emphasizes the need for more microgeographic studies of parasite-vector compatibility to more fully explore catalog the parasite and vector genes and variants that determine vector-parasite compatibility.

## Materials and Methods

### Mosquito Collection and Sample Preparation

*Anopheles* mosquitoes were collected by human landing catch following the recommendations of the National Ethical Committee in Ndenguang, Cameroon (CNERSHN° 2020/07/1259/CE/CNERSH/SP), and in Benguia, Gabon (0031/2014/SG/CNE), over the course of four missions per site between Nov 2019 and July 2021. After live capture, mosquitoes were isolated in cages for at least 10 days to allow sporozoites to develop and migrate to the salivary glands. After death, whole body DNA extraction was performed following the protocol described by Morlais et al., (2004)^47^ If mosquitoes died less than 3 days after capture, the abdomen was dissected and DNA was extracted from head and thorax to minimize detection of parasites restricted to the blood meal or midgut. Parasite infection status was determined by qPCR, targeting the 18S rRNA gene following Mangold et al. (2005)^48^ with modifications. Reactions were performed in a final volume of 10 µl containing 1× EvaGreen qPCR mix (Euromedex), 0.6 µM of each primer, and 1 µl of template DNA. Amplification consisted of an initial denaturation at 95 °C for 10 min, followed by 40 cycles of 95 °C for 10 s, 50 °C for 5 s, and 72 °C for 20 s. Melting curve analysis was performed after amplification to assess the specificity of the PCR products. Selective whole genome amplification (sWGA) was then performed two times one each DNA sample as previously described using the complete primer set pfset1^49^ to enrich parasite DNA.

### Whole Genome and Targeted Sequencing Data Generation

After sWGA, sample libraries were constructed for whole genome sequencing (WGS) using a NEBNext kit (New England Biolabs) and sequenced on an Illumina NovaSeq instrument at the Broad Institute using 150 bp paired-end reads. Sequencing of replicate sWGA products produced low read coverage for most samples, so targeted amplicon sequencing using the 4CAST panel was also performed.^50^ This panel contains amplicons capturing highly heterozygous regions of the *P. falciparum* loci *CSP, AMA1, SERA2*, and *TRAP*. Each first-round reaction contained 1.5 μL 4CAST primer mix (each at a concentration of 6.25 μM), 5 μL KAPA HiFi HotStart ReadyMix (2x), and 4 μL DNA template.

To maximize sequencing coverage of the amplicon panel markers, 2 μL of product from each sWGA product was pooled to provide the input DNA for each sample. The second round of was then conducted, each reaction consisting of 2.2 μL of unique dual index (10 μM Illumina Nextera UD indices), 5 μL KAPA HiFi HotStart ReadyMix (2x), 2 μL nuclease-free water, and 3 μL of PCR1 product. PCR2 products were pooled in equal volumes and size selection was performed using Agencourt AmpureXP beads (Beckman Coulter). Library composition and concentration were confirmed using the Agilent Bioanalyzer 2100 and qPCR (Roche) before sequencing at 4 pM with 20% PhiX in paired-end, 500 cycle format using a MiSeq Reagent Kit v2.

Three sets of PCR primers amplifying different regions of *Pfs47* were tested in a laboratory setting following parasite sWGA, using whole body DNA from NF54-infected *Anopheles gambiae* (PEST colony). Amplification regions were selected to focus on Domains 2 and 3, given those domains were previously noted to contain Single Nucleotide Polymorphisms (SNPs) potentially under selection.^22,38^ Uninfected *An. gambiae* PEST specimens were used as a negative control. Amplifications were carried out using the Qiagen Multiplex PCR kit according to manufacturer instructions. One set of primers successfully amplified *Pfs47*, spanning portions of Domains 2 and 3 (coding sequence nucleotides 672-1172), and was used for amplification of *Pfs47* in all samples: (5’AGAATCTGGAAATTGTGAAGA3’, 5’TCTGGTTTTGTTGGTAAGGCT3’). For each sample, 2 μl of DNA were pooled from each of the two independent sWGA reactions and *Pfs47* was amplified using Qiagen Multiplex PCR. The products were cleaned using the ExoSap protocol according to manufacturer instructions (ThermoFisher), and all samples were analyzed by gel electrophoresis for a band of ∼500 bp. Samples with a correctly sized band were quantified on a Qubit 4 Fluorometer and sent to Azenta Genewiz for Sanger (capillary) sequencing with both the forward and reverse primers.

### Sequencing Data Analysis

The complete mitochondrial genomes of *An. gambiae,*^51^ *An. moucheti*^52^ and *An. funestus*^53^ were aligned in MEGA11 (11.01.03) using ClustalW and the alignment was used to generate an unrooted Maximum Likelihood phylogenetic tree using ModelFinder^54^ and IQTree^55^ using the CIPRES Science Gateway. The tree was visualized in R (4.4.1) using the ape (5.8-1) in RStudio (2024.04.2+764) using tidyverse (2.0.0).

Haplotype data of *Pfs47* from Pf7^56^ were downloaded from Pf-Haploatlas^57^ to characterize global population diversity. Position-level data was extracted from the haplotypes and SNPs present in fewer than five samples were excluded.

Targeted Illumina sequencing data for the 4CAST PCR multiplex were filtered for errors using a previously described pipeline (https://github.com/broadinstitute/malaria-amplicon-pipeline)^50^ that incorporates the DADA2 de-noising algorithm.^58^ The error-filtered targeted amplicon sequencing data were analyzed in R (4.4.1) tidyverse (2.0.0). Amplicon sequence variants (ASVs) were filtered to exclude variants supported by fewer than 5 reads. Observed ASVs were also excluded when they represented less than 1% of the total read count of a given sample or were never observed as the major variant in any sample. Complexity of infection (COI; the number of genotypically distinct parasite strains per sample) was calculated for each sample as the maximum number of genotypes at a single locus, and samples were classified as monoclonal if they exhibited only one genotype per locus at all loci.

Our Sanger capillary sequencing data were analyzed in R (4.4.1) using tidyverse (2.0.0). Trace files were read and aligned to the 3D7 *Pfs47* sequence (PF3D7_1346800) as a reference using sangerseqR (1.40.0) and Biostrings (2.72.1) locally and all discrepancies were recorded. Secondary peaks indicating possible heterozygosity in multiclonal infections were recorded if they exhibited a height greater than 33% the height of the primary peak. SNPs were first filtered for agreement between the forward and reverse reads to identify the most common variants. For nucleotide positions of interest (coding sequence coordinates 742, 814, 815, 1106) both forward and reverse reads were recorded if there was discrepancy between them. Analysis and visualization were performed in RStudio (2024.04.2+764) using tidyverse (2.0.0) and ggplot2 (3.5.1), along with Hmisc (5.2-1) to calculate confidence intervals for plotting, ggpubR (0.6.0) to arrange multiple plots, exact2x2 (1.6.9) to run Fisher’s exact tests, rehh (3.2.2) to generate bifurcation diagrams, sjPlot (2.9.0) to plot binomial logistic regression models, and lubridate (1.9.4). All annotated scripts used for data analysis are available on Github.

### AlphaFold structure of *Pfs47* + Pymol

The predicted structure of *Pfs47* was downloaded from AlphafoldDB (AF-Q8IDN0-F1-v6) and visualized in Pymol (3.0.3). All three domains were predicted with high confidence (Qscore > 90%). For clarity, the GPI anchor (residues 413-439) and signal peptide (1-27) were removed.

## Acknowledgments

This work was supported by NIH grant U19AI110818 (D.E.N.), and by ANR-18-CE35-0002-01–WILDING (D.A.).

