## Supplemental Information for "Genotypic evidence of vector-mediated selection on *Plasmodium falciparum* in infected field-collected *Anopheles* mosquitoes"

^+^Co-second authors

*Daniel E. Neafsey

**This PDF file includes:**

Figures S1 to S4

Table S1

Figures

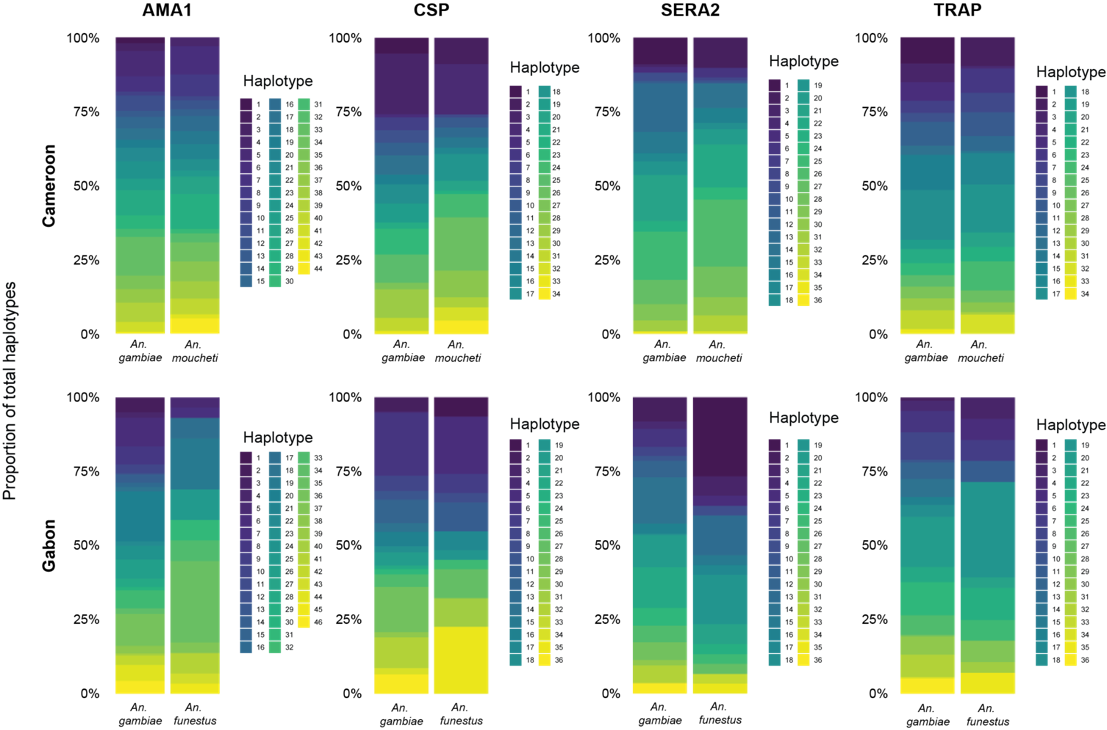

**Fig. S1.** **Haplotype distributions from 4CAST Amplicon sequencing**

Distribution of the prevalence of haplotypes for each amplicon sequenced in 4CAST. Haplotypes are numbered by overall prevalence.

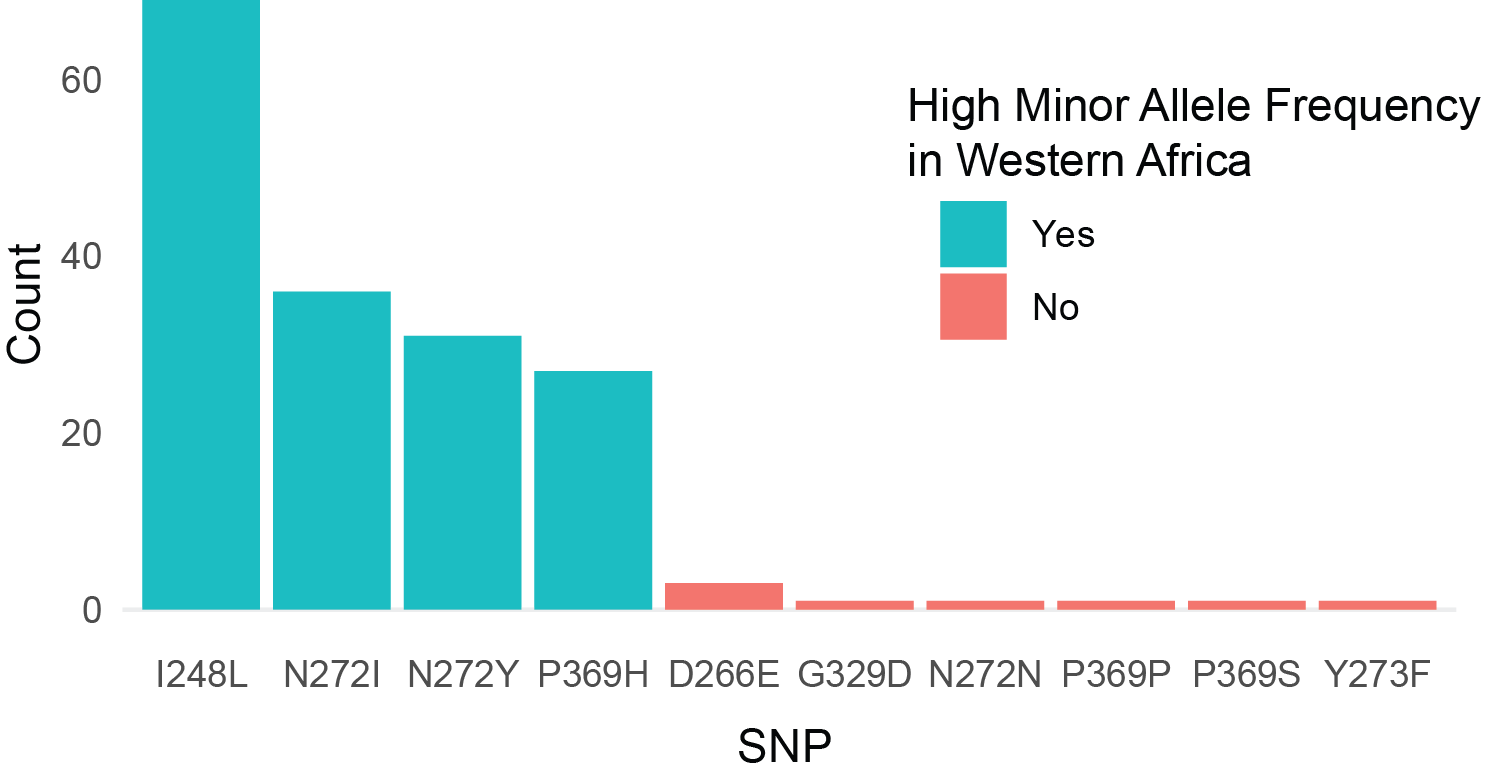
**Fig. S2.** **Frequency of each *Pfs47* SNP in our dataset**

Total count of each SNP that appeared in our sequencing results after filtering for agreement between forward and reverse reads. SNPs that had high prevalence in the Pf7 dataset are marked in blue.

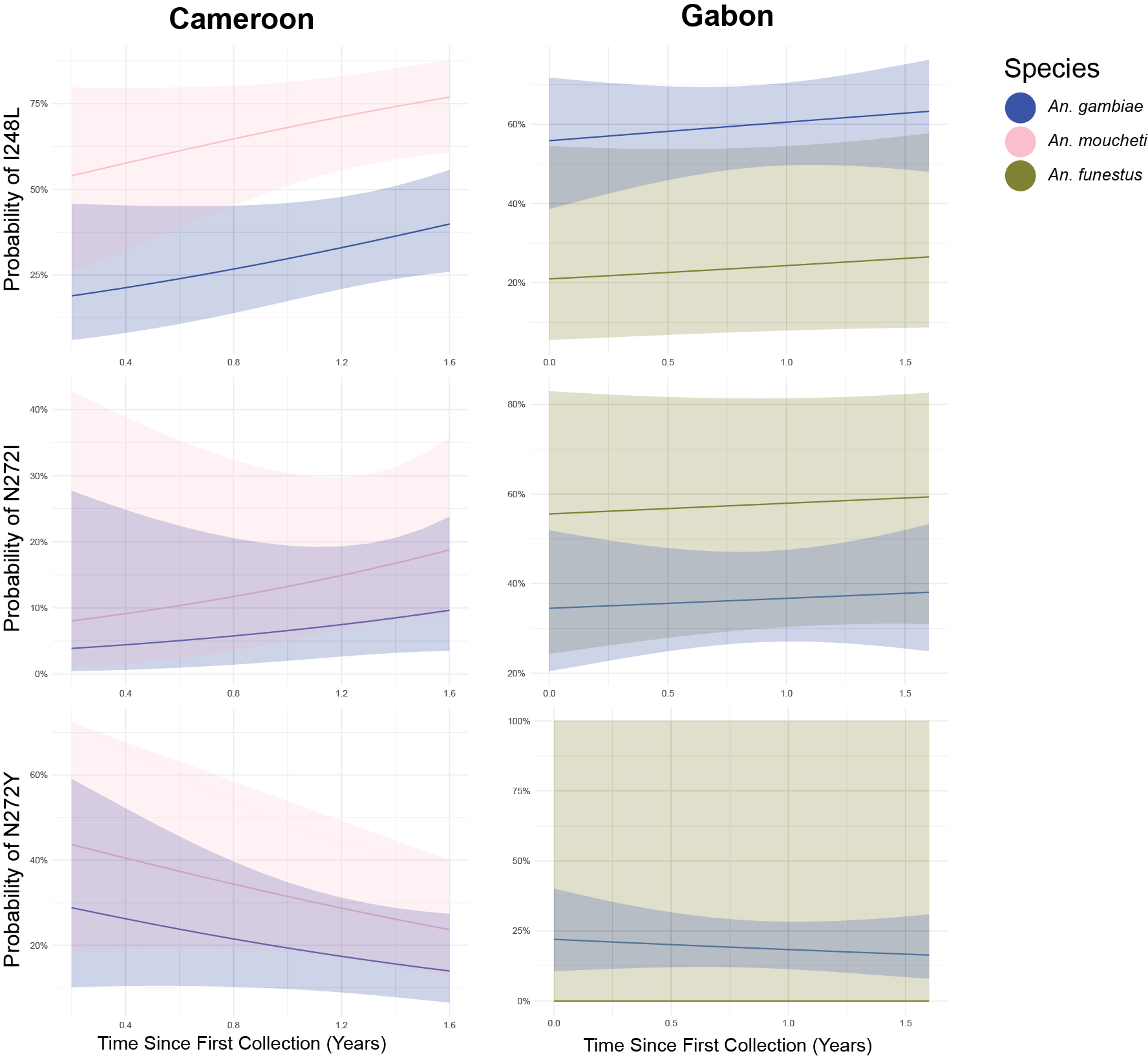

**Fig. S3.** **Binomial Logistic Regression Models for SNP Prevalence**

Logistic regression model results plotted for each SNP at each collection location. Both species and date were added as predictor variables. Standard Error is plotted as a shaded area.

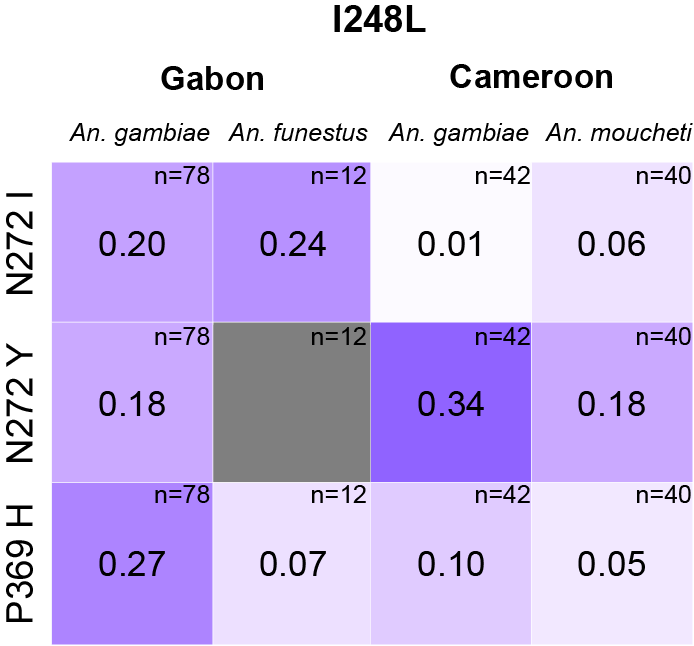

**Fig. S4. R-squared values for each residue against I248L**

R-squared values were generated by finding the coefficient of linkage disequilibrium (D) for N272I, N272Y, and P369H paired with I248L.

Tables

Table S1. Mosquito Collection Extended Information

| **Country** | ***Anopheles* Species** | **Mission** | **Collected** | **Infected** | ***Plasmodium* Species** | | | | ***P. falciparum* Prevalence** |
| --- | --- | --- | --- | --- | --- | --- | --- | --- | --- |
|  |  |  |  |  | *P. falciparum* | | *P. malariae* | *P. ovale* |  |
| **Cameroon** | *An. funestus* | M3 | 3 | 0 | 0 | 0 | | 0 | **0.000** |
|  |  | M4 | 1 | 0 | 0 | 0 | | 0 |  |
|  | *An. gambiae* | M1 | 473 | 24 | 18 | 5 | | 1 | **0.064** |
|  |  | M2 | 489 | 28 | 26 | 1 | | 1 |  |
|  |  | M3 | 123 | 0 | 0 | 0 | | 0 |  |
|  |  | M4 | 267 | 46 | 42 | 3 | | 1 |  |
|  | *An. hancocki* | M2 | 5 | 1 | 1 | 0 | | 0 | **0.200** |
|  | *An. moucheti* | M1 | 904 | 45 | 31 | 11 | | 3 | **0.041** |
|  |  | M2 | 256 | 5 | 5 | 0 | | 0 |  |
|  |  | M3 | 54 | 3 | 3 | 0 | | 0 |  |
|  |  | M4 | 827 | 55 | 44 | 7 | | 4 |  |
|  | *An. paludis* | M1 | 12 | 0 | 0 | 0 | | 0 | **0.000** |
|  |  | M2 | 180 | 0 | 0 | 0 | | 0 |  |
|  |  | M3 | 9 | 0 | 0 | 0 | | 0 |  |
|  |  | M4 | 68 | 0 | 0 | 0 | | 0 |  |
|  | *An. ziemanni* | M2 | 178 | 0 | 0 | 0 | | 0 | **0.000** |
|  |  | M3 | 7 | 0 | 0 | 0 | | 0 |  |
|  |  | M4 | 6 | 0 | 0 | 0 | | 0 |  |
|  | sp | M1 | 4 | 0 | 0 | 0 | | 0 | **0.000** |
| **Gabon** | *An. coustani* | M1 | 25 | 0 | 0 | 0 | | 0 | **0.000** |
|  |  | M2 | 10 | 0 | 0 | 0 | | 0 |  |
|  |  | M4 | 7 | 0 | 0 | 0 | | 0 |  |
|  | *An. coustani/ An. ziemanni* | M1 | 4 | 0 | 0 | 0 | | 0 | **0.000** |
|  |  | M3 | 118 | 0 | 0 | 0 | | 0 |  |
|  |  | M4 | 9 | 0 | 0 | 0 | | 0 |  |
|  | *An. demeilloni* | M1 | 2 | 0 | 0 | 0 | | 0 | **0.000** |
|  | *An. dureni* | M1 | 1 | 0 | 0 | 0 | | 0 | **0.000** |
|  | *An. funestus* | M1 | 101 | 2 | 2 | 0 | | 0 | **0.048** |
|  |  | M2 | 48 | 1 | 1 | 0 | | 0 |  |
|  |  | M3 | 56 | 4 | 4 | 0 | | 0 |  |
|  |  | M4 | 151 | 14 | 10 | 3 | | 1 |  |
|  | *An. funestus/ An. nili* | M1 | 2 | 0 | 0 | 0 | | 0 | **0.000** |
|  | *An. gambiae* | M1 | 1271 | 65 | 56 | 6 | | 3 | **0.064** |
|  |  | M2 | 303 | 11 | 11 | 0 | | 0 |  |
|  |  | M3 | 196 | 23 | 19 | 2 | | 2 |  |
|  |  | M4 | 373 | 65 | 52 | 11 | | 2 |  |
|  | *An. gambiae/ An. garnhami* | M1 | 1 | 0 | 0 | 0 | | 0 | **0.000** |
|  | *An. garnhami* | M1 | 1 | 0 | 0 | 0 | | 0 | **0.000** |
|  | *An. hancocki/ An. brohieri* | M1 | 6 | 0 | 0 | 0 | | 0 | **0.000** |
|  |  | M2 | 7 | 0 | 0 | 0 | | 0 |  |
|  |  | M3 | 4 | 0 | 0 | 0 | | 0 |  |
|  | *An. moucheti* | M1 | 7 | 0 | 0 | 0 | | 0 | **0.000** |
|  |  | M4 | 7 | 0 | 0 | 0 | | 0 |  |
|  | *An. paludis* | M1 | 68 | 0 | 0 | 0 | | 0 | **0.002** |
|  |  | M2 | 191 | 0 | 0 | 0 | | 0 |  |
|  |  | M4 | 291 | 1 | 1 | 0 | | 0 |  |
|  | *An. paludis/ An. coustani* | M1 | 1 | 0 | 0 | 0 | | 0 | **0.000** |
|  | *An. paludis/ An. tenebrosus* | M1 | 1 | 0 | 0 | 0 | | 0 | **0.000** |
|  | *An. rufipes* | M1 | 1 | 0 | 0 | 0 | | 0 | **0.000** |
|  | *An. seydeli/ An. brunnipes* | M1 | 1 | 0 | 0 | 0 | | 0 | **0.000** |
|  | *An. squamosus/ An. cydippis* | M1 | 3 | 0 | 0 | 0 | | 0 | **0.000** |
|  | *An. squamosus/ An. cydippis* | M3 | 2 | 0 | 0 | 0 | | 0 | **0.000** |
|  | *An. tenebrosus* | M1 | 7 | 0 | 0 | 0 | | 0 | **0.000** |
|  | *An. tenebrosus/ An. ziemanni* | M1 | 1 | 0 | 0 | 0 | | 0 | **0.000** |
|  | *An. ziemanni* | M1 | 14 | 0 | 0 | 0 | | 0 | **0.000** |
|  |  | M2 | 1 | 0 | 0 | 0 | | 0 |  |
|  |  | M4 | 13 | 0 | 0 | 0 | | 0 |  |
|  | *An. ziemanni/ An. nami* | M1 | 6 | 0 | 0 | 0 | | 0 | **0.000** |
|  | sp | M1 | 140 | 10 | 8 | 1 | | 1 | **0.055** |
|  |  | M2 | 20 | 0 | 0 | 0 | | 0 |  |
|  |  | M3 | 40 | 4 | 3 | 1 | | 0 |  |
|  |  | M4 | 37 | 4 | 2 | 2 | | 0 |  |
